# Failure to engage the TPJ-pSTS during naturalistic scene processing in schizophrenia

**DOI:** 10.1101/828988

**Authors:** Gaurav H. Patel, Sophie C. Arkin, Daniel Ruiz-Betancourt, Fabiola I. Plaza, Safia A. Mirza, Daniel J. Vieira, Nicole E. Strauss, Casimir C. Klim, Juan P. Sanchez-Peña, Laura P. Bartel, Jack Grinband, Antigona Martinez, Rebecca A. Berman, Kevin N. Ochsner, David A. Leopold, Daniel C. Javitt

## Abstract

The ability to search for and detect social cues, such as facial expressions of emotion, is critical to the understanding of complex dynamic social situations. This ability involves the coordinated actions of multiple cognitive domains, including face-emotion processing, mentalization, and visual attention. Individuals with schizophrenia are generally impaired in social cognition, and have been shown to have deficits in all of these domains. However, the whether the neural substrates of these impairments are shared or separate remains unclear. One candidate region for a shared substrate is the right temporoparietal junction/posterior superior temporal sulcus (TPJ-pSTS), which contains areas belonging to all of the cortical networks underlying these domains.

Here we use functional MRI to examine differences in cortical activity evoked by a naturalistic movie, and link these results to impaired visual scanning and social cognition. 27 schizophrenia participants and 21 healthy controls watched a 15-minute clip of the movie “The Good, the Bad, and the Ugly” while high resolution multiband BOLD-fMRI activity was recorded. Inter-subject correlation was used to measure the evoked activity. BOLD-fMRI activity was also correlated with motion content in the movie, with the average activity in other cortical areas, and with frequency of saccades made during the movie. Visual scanning performance was measured in a separate behavioral experiment, and social cognition measured by The Awareness of Social Inference Test (TASIT).

Contrasting the groups revealed that the TPJ-pSTS has the largest engagement deficit in both cortical hemispheres in schizophrenia patients versus healthy controls. Follow-up analyses find that brain activity in this region is less correlated with the motion content of the movie, that this region is abnormally synchronized to the other cortical areas involved in the cognitive domains underlying visual scanning of social scenes, and that activity this region is less correlated with the saccades made during the movie. Lastly, schizophrenia participant visual scanning performance of this clip was impaired compared to healthy controls, and correlated across the two groups with social cognition.

These results indicate that the TPJ-pSTS plays less of an integral role in the coordination of face-emotion processing, mentalization, and visual attention in schizophrenia participants versus healthy controls. This functional deficit then impacts the visual scanning of a complex dynamic visual scenes, which in turn affects the comprehension of that scene. These findings indicating that the TPJ-pSTS is potentially the shared substrate for all of these deficits will lead to new treatments targeting this region to improve social cognition in individuals with schizophrenia.

## Introduction

Deficits in social cognition are a major source of disability in schizophrenia (1). Social cognition is a broad term that encompasses a number of domains (2). Two of these domains are the visual scanning for and the processing of social cues (such as facial expressions of emotion), and a third is mentalizing, also known as theory of mind. These domains are thought to be intimately related as the detection and processing of social cues is critical to understanding the mental states of individuals in a social situation (3). While all three of these domains are thought to be impaired in SzP, whether there is one common deficit or multiple deficits in the underlying neural substrates remains unclear (1). Here, we examine the simultaneous activation of the neural substrates underlying all three domains to determine where deficits may be located.

The neural substrates of these three domains span the cortex: face emotion recognition involves a network of areas in occipital and temporal lobes (4), visual scanning or attention involves a network of areas in parietal and prefrontal cortex (3), and mentalization involves a network of areas temporal, parietal, and both medial and lateral prefrontal cortex that is heavily overlapping with the default mode network (5). Previous studies in SzP have found potential deficits distributed throughout all of these networks, but thus far no one common area or region has emerged.

One candidate for a common region is the temporoparietal junction/posterior superior temporal sulcus (TPJ-pSTS). The cortical networks underlying all three domains have nodes in the TPJ-pSTS (6). The combination of all of these systems in a single region makes it a hub for cognitive operations (7). According to our proposed model, the TPJp-pSTS receives input about face emotion and other biological motion from the pSTS and compares it to internally maintained models in the posterior TPJ (TPJp) about the mental states of individuals in a social scene. The mismatch of the incoming sensory information and the expectation generated by the model (prediction error) then activates the anterior portion of the TPJ (TPJa), a node in the ventral attention network, causing attention and visual gaze to be reoriented to the relevant visual feature.

The arrangement of these areas within the TPJ-pSTS may be unique to humans, allowing for the quick visual scanning of the dynamic and complex social situations that humans have to navigate (6). While previous studies have noted deficits in SzP in this region for face-emotion recognition, attention, and mentalizing, these studies have typically used tasks designed to isolate the functioning of only one of these domains at a time, leaving open the question of how and where the deficits may inter-relate. We hypothesize that the common deficit lies within the TPJ-pSTS, which then would have an impact on all of the systems linked to this hub.

While the highly controlled stimuli typically used in experiments are good at isolating and examining the functioning of specific areas or cognitive domains, naturalistic experiments are able to examine the relative contributions of multiple domains at once. However, they can be difficult to analyze and interpret. One of the difficulties is the inability to tie any known sensory or cognitive operations to the patterns of activity generated by the stimuli. However, recent advances in computer vision allow for the automated classification of visual features, which can then be correlated with neural activity (8; 9). Another difficulty of using naturalistic imaging to study that is specific to the TPJ-pSTS is its high degree of morphological variability, which combined with its human-unique structure makes it difficult to resolve and separate the multiple adjacent nodes of cortical systems that lie within the TPJ-pSTS. Yet another difficulty lies in understanding the directionality of the deficits—since there is no temporal control over the presentation of stimuli, it can be difficult to understand how information flows through the many areas activated by the naturalistic stimuli.

In this study, we compare activation of cortex by a naturalistic cinematic movie between SzP and demographically matched HCs. We developed and adapted analytic tools to characterize brain activity associated with both complex movie context and patterns of eye-movements. Applying these methods to high-resolution functional imaging sequences and functional localizers to separate adjacent regions of interest (ROIs) in the TPJ-pSTS, we reveal new deficits not previously described in SzP. We first identify cortical regions showing the largest difference between groups using intersubject correlation (ISC), which compares the time-locked activation of each ROI between individuals without making assumptions of what aspect of the movie or which ROIs may be driving the differences. We then examine how TPJ-pSTS activity communicates or synchronizes with other ROIs involved with the transformation of visual features into a visual scanning plan, such as those involved in motion processing (middle temporal area, or MT), face processing (fusiform face area, or FFA), guidance of saccades (frontal eye-fields, or FEF), and prefrontal areas involved in cognitive control and mentalization. We then use continuous measures of visual features and eye-tracking to assess deficits within the TPJ-pSTS and elsewhere. Lastly, we show that eye-tracking performance is relevant to standardized measures of social cognition. Importantly, while the central hypothesis and focus of this manuscript is on the TPJ-pSTS, the methods used are data-driven and the results not biased to hypotheses about the TPJ-pSTS.

## Materials and Methods

### Participants

48 schizophrenia patients (SzP) and 48 healthy controls (HC) were recruited with informed consent in accordance with New York State Psychiatric Institute’s Institutional Review Board (IRB). 27 SzP and 21 HC of these participants completed the movie-watching MRI portion of the study. An overlapping group of 21 SzP and 23 HC completed the behavioral portion of the study; form this dataset one SzP and one HC had to be excluded because of blink-related artifacts in the eye-traces, leaving 20 SzP and 22 HC.

### Inclusion/Exclusion Criteria for Participants

Participants were primary English speakers, ages 18-55, devoid of history of head trauma and neurological disorders, and had normal or corrected visual acuity (20/40 or better). Participants with contraindication for MRI (artificial implants and/or claustrophobia) were excluded from the study. Patients met DSM-IV criteria for schizophrenia, as confirmed by the Lieber Schizophrenia Research Clinic (LSRC) before recruitment and participation. Patients were on disease-appropriate medication used within appropriate guidelines, and were assessed with the Positive and Negative Symptoms Scale (PANSS; (10)). Patients scoring above 120 on the PANSS, or those who were acutely psychotic were excluded. HC were recruited through the LSRC and through IRB-approved flyers and internet advertisements. The Structural Clinical Interview for the DSM-IV (SCID-NP) was used to exclude past or present Axis I or II disorders, significant substance use disorders in the past 6 months, and in HC significant psychiatric family history (11).

### Behavioral Session and Analyses

In the behavioral session, participants seated in front of a computer monitor with their heads resting comfortably in a head-holder. The participants first completed The Awareness of Social Inference Test (TASIT) (12) with eye-tracking. They then were instructed to watch a video clip of the first 15 minutes of the cinematic movie “The Good, the Bad, and the Ugly” (United Artists, 1966) with the sound removed and with eye-tracking, and to briefly summarize the plot afterwards to ensure attention.

TASIT performance was scored for percentage correct out of 64 total questions across 16 video clips; this data overlaps the data used in (13). Eye-tracking performance was summarized by first z-transforming the distance for each individual’s eye-position versus the mean of the HC population (distance/std. dev.) for each video frame, and then averaging across video frames to calculate an average z-transformed distance for each individual.

### MRI Session

Structural T1 and T2 (0.8mm isotropic), multiband (MB) fMRI (2mm isotropic, TR=850ms, MB factor 6), and distortion correction scans (B0 fieldmaps) were acquired as required for use of the Human Connectome Project (HCP) processing pipelines. For the movie-watching scan, subjects viewed the same cinematic movie clip used in the behavioral session, without sound and with eye-tracking, and the BOLD data was collected as one continuous 15 minute acquisition (1049 MR frames).

### Image Processing

MRI data were preprocessed using the Human Connectome Processing pipelines v3.4 (14), which places the data into grayordinates in a standardized surface atlas (as opposed to voxels in a volume atlas). The functional data were additionally cleaned of artifact largely following the recommendations from Power *et al.* (15). To equalize numbers of censored frames, we used an adaptive framewise displacement (FD) threshold that set the threshold as the 75%tile + 1/2 * interquartile range of the FD trace for each run, limited to a range between 0.2mm and 0.5mm. Numbers of frames censored did not differ significantly (HC=184.4(37.9), SzP=230.2(34.1), t_46_=1.85, p=0.07) and all results reported below were similar when analyses were repeated with a universal threshold of FD=0.2mm.

### Intersubject Correlation Analyses

TPJ-pSTS regions of interest were derived from localizer tasks (described in **Supplemental Materials**). For intersubject correlation (ISC) analyses, we correlated the time-course of BOLD activity of for each grayordinate or ROI in each individual with each of the HC individuals.

### TPJ Synchronization with Other ROIs

We next examined group differences in the synchronization of TPJ ROIs with the movie-evoked activity in ROIs from the other cortical areas involved in transforming visual inputs into eye-movements. This measure, similar to the intersubject functional correlation described in Simony *et al.* (16), reveals deficits in the stimulus-driven synchronization of activity between ROIs in SzP. The average HC time-course for each ROI was used as the reference to negate the impact of processing deficits *within* these other ROIs in SzP affecting the measurement. This measure, unlike typical ROI-ROI functional connectivity measures, is also directional—for instance, HC V1→ SzP V2 synchronization may be decreased compared to HC V1→HC V2 synchronization, but that does not necessitate that HC V2→ SzP V1 synchronization is also decreased compared to HC V2→HC V1. For this measure, we first extracted the average time-course of activity for each non-TPJ ROI (R MT, R FFA, R FEF) from the average of all of the HCs. This time-course was then correlated to the mean time-course of each TPJ ROI (TPJp, TPJm, and TPJp) for each individual. Grayordinate maps were created for visualization by correlating the HC ROI time-courses with all grayordinate time-courses. In exploratory analyses, we also examined TPJ synchronization with other cortical ROIs involved in social cognition.

### Visual Feature and Saccade Correlations with BOLD Activity

Finally, we examined the correlation of BOLD activity with continuous measures of the intensity of visual features in the movie as well the number of saccades per MR frame (saccade frequency). Low-level visual features (motion, contrast, luminance) were extracted from the movie as described in Russ *et al.* (8) with the exception of log-transformation of the motion parameter for normalization. Automated face detection (http://aws.amazon.com/rekognition) was used to score each MR frame for the number of faces. Automated saccade detection (SR Research, Mississauga, Ontario, Canada) was used to score each MR frame for the number of saccades. These regressors were convolved with a hemodynamic response function (17) and then correlated to the time-course of activity for each ROI or grayordinate.

### Group Comparison Statistics

For all relevant analyses, HCs were compared to leave-one-out averages of all other HCs. Correlation values were Fisher-z transformed, and then compared between groups using repeated measures ANOVAs and post-hoc t-tests. All values are reported as mean(confidence interval). Grayordinate-wise ISC contrast was calculated using mixed-effects in FEAT (18) and cluster inference threshold set at p<.05 by PALM (19).

## Results

### Demographics and MRI Eye-tracking Statistics

SzP and HC were demographically similar in age, gender, race/ethnicity, education, socioeconomic status (SES), handedness, and IQ (**Table 1**). SzP were mostly mildly to moderately ill (PANSS=58.4(14.9)) on antipsychotic medication (mean CPZ dose=691.9(1034.5) mg).

In the 14 SzP and 13 HC with usable eye-tracking in the MRI, mean number of video frames dropped due to artifact (such as blinks) was similar (30.9(20.7)% versus 22.9(17.8)%, t_25_=1.1, p=0.3). The mean number of saccades performed during the free-viewing of the movie in the MRI was also similar in SzP versus HC (1646.8(542.7) versus 1666.5(589.7), t_25_=0.09, p=0.92).

### Intersubject Correlation

We first investigated whether there were brain areas that responded differently to the movie between SzP and HC, using ISC to quantify engagement. Qualitatively, overall ISC patterns were similar in the two populations, with extensive engagement of visual cortex in occipital and temporal lobes, as well was sections of parietal and both lateral and medial prefrontal cortex (**Figure 1A** and **B**). However, contrasting the ISC patterns in the two populations revealed a focal ISC deficit in the TPJ (**Figure 1C**). We did not observe any other large ISC deficits in any other cortical region in either hemisphere.

**Figure 1:**
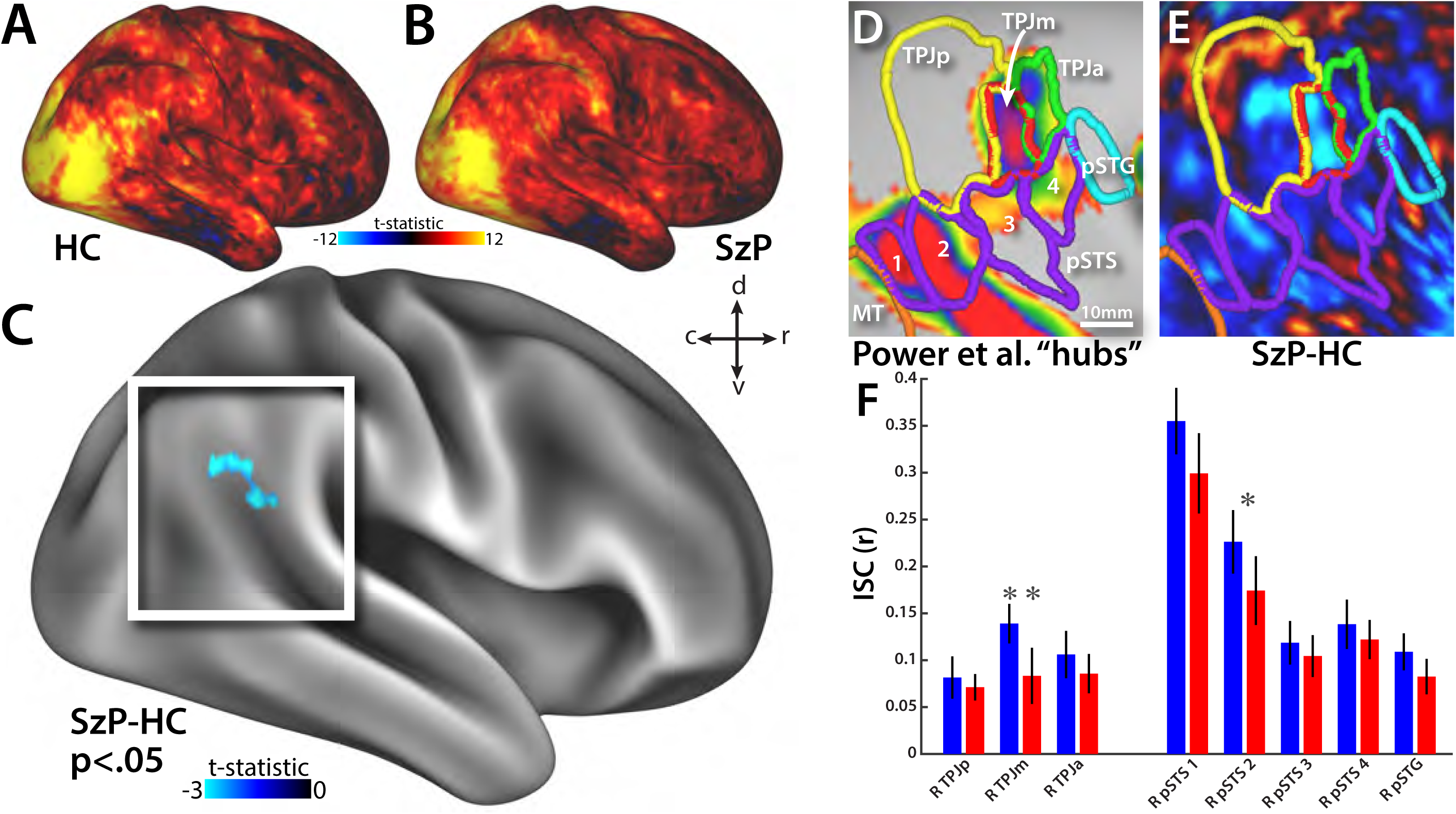
Grayordinate-wise comparison of ISC reveals focal TPJ-pSTS deficits. Right hemisphere views of HC (**A**) and SzP (**B**) ISC maps demonstrate strong engagement of occipitotemporal visual, dorsal attention, pSTS, and prefrontal areas. Contrast map reveals large focal deficit that is the only focus to survive cluster correction (**C**). No cluster survives comparison in the left hemisphere. White frame in **C** outlines region shown in **D** and **E. D)** TPJ-pSTS subdivided into mentalization (TPJp, yellow), ventral attention (TPJa, green), face-emotion processing (pSTS, purple), and dorsal attention (pSTG, cyan) ROIs. Central TPJ ROI (TPJm, red) falls within a hub defined by Power *et al.* (7). **E)** Focal ISC deficit falls mostly within TPJm. **F)** ROI by ROI group comparison of ISC in the TPJ-pSTS. *p<.05, **p<.01.

To localize the deficits within the TPJ-pSTS, we used task localizers to subdivide the TPJ-pSTS ISC activations into functionally defined regions of interest (ROIs) (**Figure 1D** and **Supplemental Figure 1**). Localized ROIs included the four ROIs in the pSTS involved in processing of moving facial expressions (purple), a mentalizing ROI in the TPJp (yellow), and a selective attention related ROI in the TPJa. An additional ROI that may be a node of the dorsal attention network (pSTG, cyan) (20) was also identified. In between these ROIs was another ROI we labeled TPJm. TPJm overlapped a right TPJ hub area defined by Power *et al.* (7) (**Figure 1D**).

The ISC deficits we observed in the SzP were largely concentrated in the TPJm (**Figure 1E**). This observation was confirmed by ROI-level statistics comparing group engagement of the 8 TPJ-pSTS ROIs (**Figure 1F**). We found a significant difference between groups (F_1,46_=6.1, p=0.016) and a significant group x ROI interaction (F_1,46_=5.7, p=0.021). In post-hoc t-tests, TPJm exhibited the most significant difference (t_46_=3.0, p=0.0048) with an additional difference in pSTS 2 (t_46_=2.1, p=0.041).

### Synchronization of TPJ activity with Visual Processing, Attention/Oculomotor, and Other Areas

We then examined whether there were deficits in the synchronization of TPJ-pSTS activity with visual processing (MT, FFA) and attention/oculomotor areas (FEF) that send information to or receive information from the TPJ-pSTS in the process of transforming visual information into a saccade plan. In both populations, MT activity was highly synchronized with other occipitotemporal visual areas, TPJ-pSTS ROIs, and frontoparietal and dorsolateral regions of the brain usually associated with selective attention and other cognitive processes (**Figure 2A** and **B**). However, contrasting these synchronization maps revealed focal deficits in the TPJ for SzP (group: F_1,46_=11.7, p=0.001) (**Figure 2C**). with a greater impact on TPJp than TPJm or TPJa (group x TPJ ROI: F_1,46_=9.1, p=0.004) (**Figure 2D**). Repeating the same analysis for R FFA (face-processing, **Figure 2E**) and R FEF (attention/oculomotor, **Figure 2F**) revealed a similar pattern of reduced synchronization with TPJ activity in SzP (group: F_1,46_=12.8/9.0, p=0.0008/0.004) with a similar difference between TPJ ROIs (group x TPJ ROI: F_1,46_=11.8/7.2, p=0.001/0.01), though in R FFA the greatest deficit was with TPJm. Synchronization with pSTS did not differ between groups for any of these areas.

**Figure 2:**
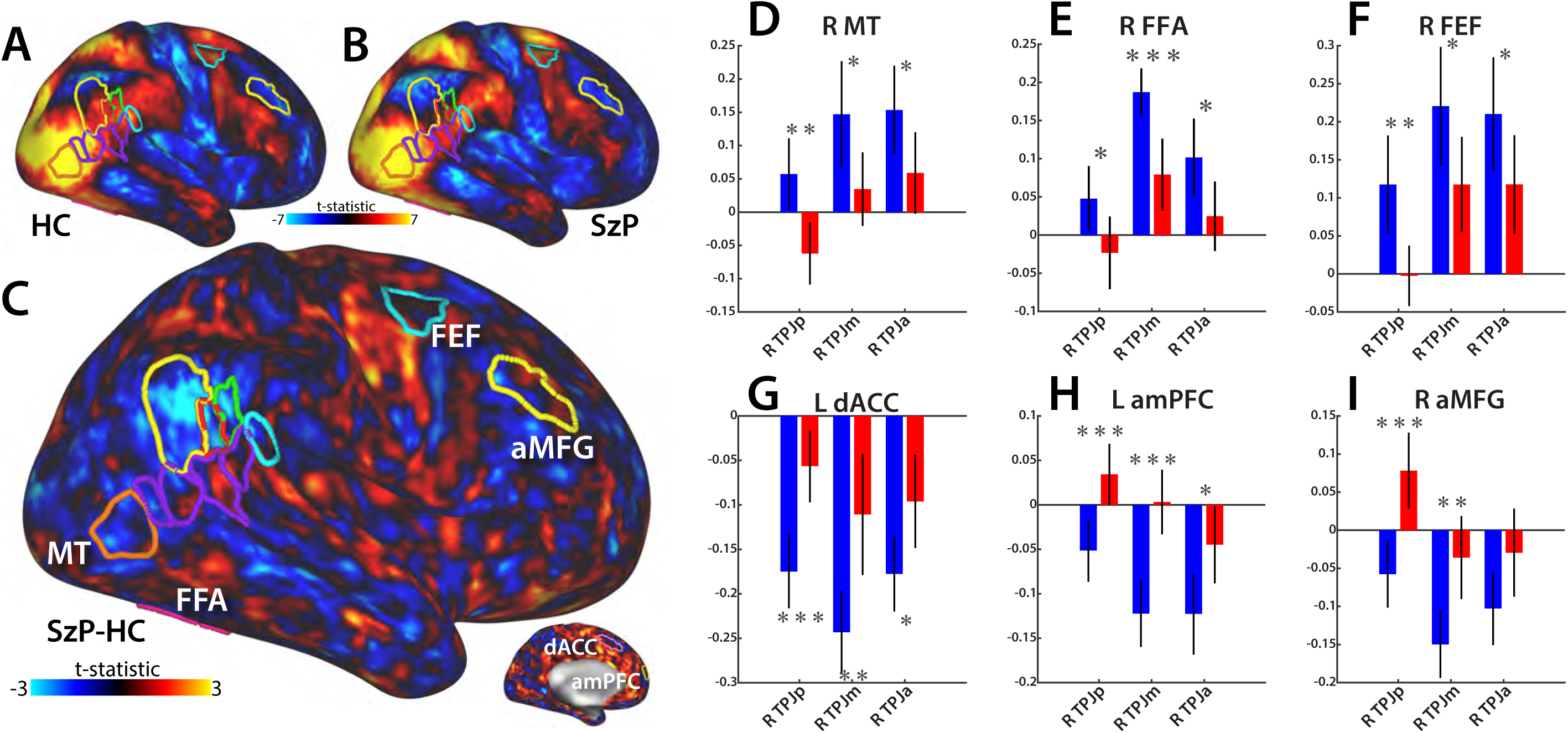
Differences in TPJ synchronization with other ROIs. **A and B)** Grayordinate synchronization maps of average HC MT activity demonstrating robust correlations in both populations with occipitotemporal visual, dorsal attention, pSTS, and prefrontal areas. **C)** Grayordinate contrast reveals large deficit centered within TPJ ROIs. **D)** MT→TPJ ROI synchronization deficits in SzP. **E and F)** Similar deficits for FFA and FEF. **G, H, and I)** Increased synchronization of medial and lateral prefrontal areas with the TPJ. *p<.05, **p<.01, ***p<.001. *MT:* middle temporal. *FFA:* fusiform face area. *FEF:* frontal eye-fields. *dACC:* dorsal anterior cingulate. *amPFC:* anteromedial prefrontal cortex. *aMFG:* anterior middle frontal gyrus.

In exploring synchronization with other cortical areas involved in social cognition, we found a pattern of SzP having decreased synchronization for all the TPJ areas involved in attention (TPJa) and mentalization (TPJp), as well as the “hub” area (TPJm), for most visual and face processing ROIs, IPS and FEF dorsal attention ROIs, and the TPJ-pSTS ROIs themselves. However, a number of prefrontal areas demonstrated the inverse pattern. These included areas within the cingulo-opercular network activated by the visual search task (e.g. L dorsal anterior cingulate cortex (L dACC) in **Figure 2G**), the default mode network activated by the mentalization localizer (e.g. L antero-medial prefrontal cortex (L amPFC) in **Figure 2H**), and the frontoparietal network also activated by the mentalization localizer (e.g. L anterior middle frontal gyrus (R aMFG) in **Figure 2I**). These results suggest that there is *increased* synchronization between the prefrontal areas and the three TPJ ROIs in SzP versus HC. Of note, when these analyses were performed without global signal regression, all values were shifted positively but maintained the same relative differences.

### Correlation of BOLD Activity with Visual Features

We next examined differences in the correlation of TPJ BOLD activity with the visual features present in the movies. In both groups, the highest correlations were observed with motion, with voxels correlated with motion speed spanning visual cortex into the TPJ-pSTS, extending beyond the cortex activated by the motion localizer and mirroring the pattern evoked by the face-emotion localizer (**Figure 3A** and **B** versus **Supplementary Figures 1E** and **1G**). BOLD activity correlations with motion also demonstrated the largest difference between groups (group: F_1,46_=8.6, p=0.005). There was also a significant group x TPJ ROI effect (group: F_1,46_=7.4, p=0.01), with TPJm demonstrating the largest deficit in SzP versus HC (t_46_=2.87, p=0.006) (**Figure 3C** and **D**). Visual contrast demonstrated only weaker effects of group (group: F_1,46_=7.4, p=0.01) but no group x TPJ ROI interaction, and neither luminance or faces demonstrated any differences. None of the visual feature-BOLD correlations differed strongly in any pSTS areas.

**Figure 3:**
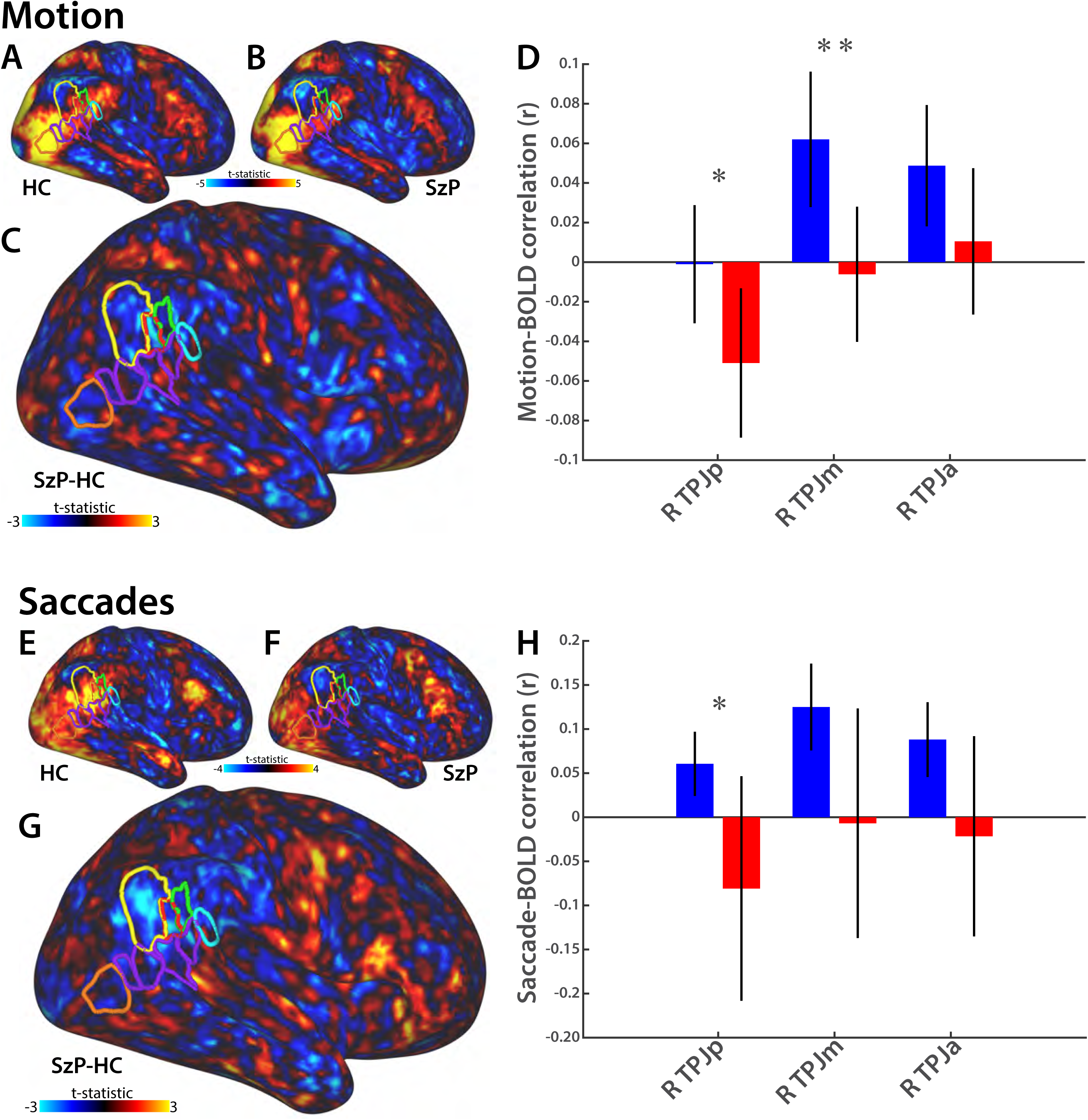
Motion and saccade frequency correlations with BOLD activity. **A and B)** Right lateral views of motion correlation maps in HC (**A**) and SzP (**B**) shows strong correlations with occipital, ventral temporal, and frontoparietal areas. **C)** Grayordinate contrast reveals focal deficit within TPJ. **D)** Motion correlation deficits strongest in TPJm. **E and F)** Right lateral views of saccade frequency correlation maps in HC (**E**) and SzP (**F**) demonstrates robust correlation with occipitotemporal visual, dorsal attention, pSTS, and prefrontal areas. **G)** Grayordinate contrast reveals focal deficit within TPJ. **H)** Strongest TPJ ROI deficits observed in TPJp. *p<.05, **p<.01.

### Correlation of BOLD Activity with Saccades

For the subset of participants with usable eye-tracking data, we next compared BOLD correlations with saccades rates between groups. In both groups there was robust correlation of saccades with the BOLD activity in visual cortex and the dorsal attention network areas involved in oculomotor planning, including the posterior IPS (pIPS) and the frontal eye-fields (FEF) (**Figure 3E**). However, SzP fail to demonstrate the same magnitude of TPJ correlation as HCs (group: F_1,25_=4.6, p=0.041) (**Figure 3F-H**). There was also a significant group x TPJ ROI interaction (group: F_1,25_=4.3, p=0.048), with the largest difference in the TPJp (t_25_=2.23, p=0.03).

### Correlation of Visual Scanning and Social Cognition

We next examined whether the visual scanning patterns of this cinematic movie predicted social cognition performance (as measured by TASIT (12)). Given the limited number of participants with usable eye-tracking data in the MRI, we instead used eye-tracking data from the participants who viewed the same movie clip in a behavioral set-up outside of the MRI. SzP versus HC were substantially impaired in TASIT performance (68.5(12) versus 85.4(6.7), t_39_=5.7, p=10^−6^). They were also impaired in visual scanning (1.48(.26) versus 1.26(.15), t_39_=3.3, p=0.002) (**Figure 4**). Visual scanning performance was strongly correlated with TASIT performance after accounting for group with no interaction of group and visual scanning (visual scanning: F_1,37_=8.3, p=0.007, group: F_1,37_=16.6, p=0.0002, group x visual scanning: F_1,37_=0.17, p=0.7). There was no correlation of TASIT performance or visual scanning with antipsychotic medication dose.

**Figure 4:**
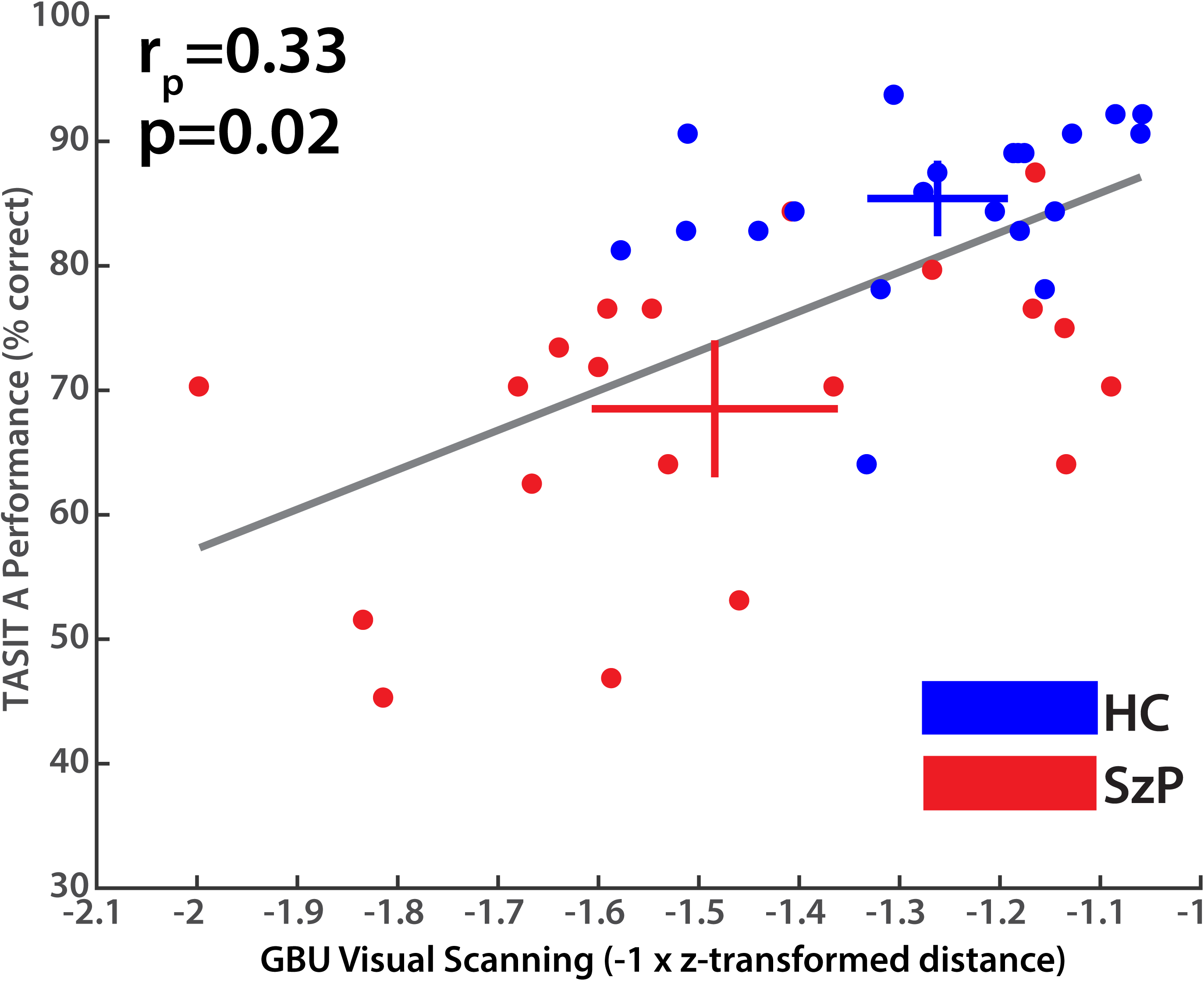
Visual scanning of “The Good, the Bad, and the Ugly” correlates with TASIT performance. Partial correlation shown after accounting for group membership correlation with TASIT performance. Cross-bars show group confidence intervals for each measure.

## Discussion

In summary, we found convergent deficits in SzP in the engagement of the TPJ-pSTS by a naturalistic movie that diminish the role this region plays in transforming visual information into a saccadic plan, which then potentially affects social cognition. Specifically, the movie failed to engage a central area within the TPJ-pSTS—TPJm—which also did not respond to appropriately to visual motion in the movie. In addition, the activity in the TPJm, TPJp, and TPJa all failed to appropriately synchronize with visual processing and attention areas and did not correlate with saccades. Lastly, we found evidence of abnormally *increased* correlation of SzP TPJ activity with cognitive control and mentalization ROIs, especially in the TPJp. Together, these results indicate that the TPJ did not play the same role in visual processing, attention, and saccade planning as it does in HCs.

Together, these results suggest that the TPJ-pSTS response to motion is abnormal in SzP. These results are in line with previous behavioral findings about reduced sensitivity to motion in general (21-23) and facial expressions and other types of biological motion more specifically (24-26). The similarity of the pattern of motion-correlated activation through the pSTS with the face-emotion localizer activity suggests that most of the motion in the movies was related to facial expression motion. However, unlike previous fMRI studies in SzP (21; 23) we do not find that the functional deficits lie within the motion/biological motion processing areas in MT or the pSTS; rather they lie in the TPJ, a region not generally implicated directly in sensory processing (6). This result may indicate that lower level motion processing deficits prevent these signals from reaching the TPJ, but that motion processing deficits in these lower level areas themselves are being obscured by other visual and cognitive signals. Indeed, the MT and pSTS signals correlate with themselves and other visual processing/attention areas similarly in HC and SzP (**Figure 2A-B**).

This motion processing failure appears to directly impact an area in the heart of the TPJ, TPJm (**Figure 5**). Another recent movie-watching study found a similar deficit, though based on the Yeo *et al.* seven network parcellation scheme it was identified as a default mode network deficit (27; 28). A number of studies have indicated that the organization of this region is complex, and is made more complex by the high degree of individual variability in its anatomy (29; 30). Power and colleagues viewed this complex organization as a sign that it may be a cognitive hub (7), which fits with our finding that the TPJm lies right between the pSTS face-emotion processing areas and both the TPJp mentalization and TPJa attention areas. Parcellation schemes based on functional connectivity have found a small patch of cortex at the same location that either had its own unique connectivity profile within the TPJ (28; 31) or was difficult to classify (32). In our localizer tasks, it was not strongly activated by either mentalization or detection, and also appeared to be equally activated by static and moving faces. This may suggest that it serves as a gatekeeper area that is activated by salient stimuli such as faces. Failure to detect or properly process motion associated with facial expressions in MT or pSTS, then, may lead to its failure to activate, thus preventing these stimuli as being registered as salient. An alternative explanation is that the TPJm itself is dysfunctional, failing to appropriately integrate motion-related information from MT and pSTS with other modalities.

**Figure 5:**
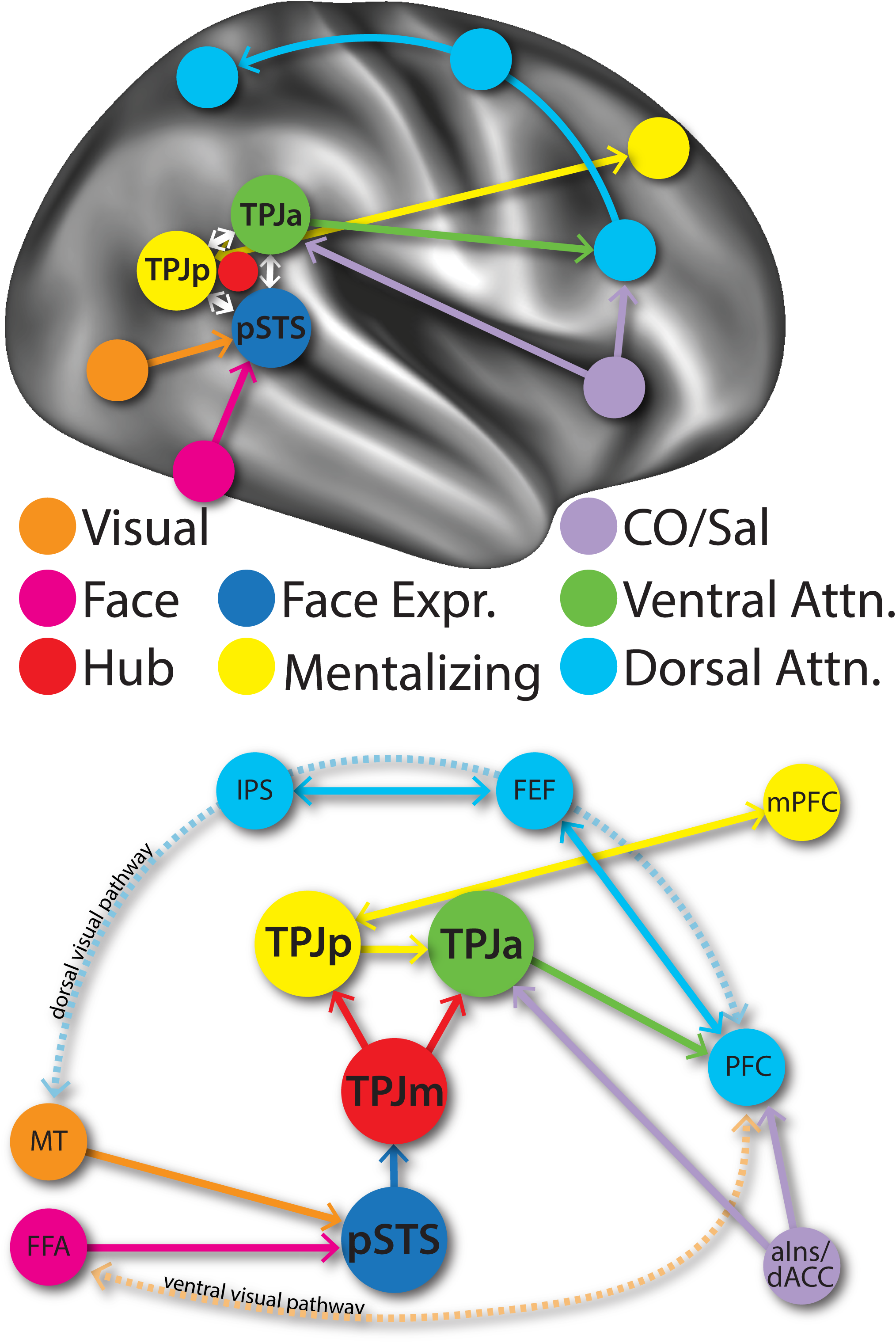
Schematic of the interactions between networks involved in visual scanning of social scenes at the TPJ-pSTS, modified from Patel *et al.* (6). The TPJ-pSTS serves as a third link between visual areas and the dorsal attention areas the control attention and saccade planning/visual scanning. Information about face motion from MT and FFA converges on the pSTS. This information is conveyed to TPJp and TPJa via TPJm. These TPJ areas are modulated by prefrontal mentalization areas, such as those in mPFC, and prefrontal cognitive control (cingulo-opercular/salience) areas, such as aIns and dACC. Based on these various inputs, the TPJp and TPJa determine whether further visual exploration of the social scene is necessary to update the ongoing mentalization operations. If further saccades are needed, the TPJa is activated, which then triggers saccade planning in the dorsal attention areas through prefrontal cortex. *MT:* middle temporal. *FFA:* fusiform face area. *IPS:* intraparietal sulcus. *FEF:* frontal eye-fields. *dACC:* dorsal anterior cingulate. *PFC:* prefrontal cortex. *mPFC:* medial prefrontal cortex. *aIns:* anterior insula. *dACC:* dorsal anterior cingulate cortex.

In any case, failure to activate the TPJm would then lead to a broader failure to engage the other TPJ areas. According to our proposed model, this would then lead to the failure to use the mentalization models maintained in the TPJp to make decisions about if and where to visually explore for additional socially relevant information, which then results in the failure to activate TPJa and to generate saccades. This failure is reflected in the decreased correlation of TPJ activity with the generation of saccades, along with the decreased synchronization of visual processing and attention area activity. This failure to engage the TPJ in saccade planning then potentially underlies the differences in the visual scanning patterns observed in the behavioral dataset, which then impacts social cognition.

While motion processing inputs may disrupt TPJ operations, our finding of increased synchronization of the TPJ with prefrontal cognitive control and mentalization areas suggest two conclusions. First, at least some of the movie-synchronized cognitive control and mentalization signals that exist in HCs watching the movie also exist in SzP. Previous studies have found deficits in mentalization in SzP (33; 34), but it has not been clear if these deficits were primary to the areas underlying mentalization or secondary to deficits in face-emotion processing (1). Our results suggest that they may be secondary to face-emotion processing deficits in the TPJ-pSTS, but this will have to be confirmed with a more direct comparison of the two processes (35).

Second, unlike the sensory input channel through the TPJm, information from these prefrontal areas is being shared with the TPJ, especially the TPJp. According to our model, this may be result from decreased visual-TPJ synchronization that biases TPJ activity to synchronize with prefrontal areas. Another explanation may be that prefrontal-TPJ synchronization may be abnormally increased, reflecting either increased connectivity of the TPJp (a node in the default mode network (36)) with other default mode network areas (37) or areas in other networks (38)(39). Disentangling these possibilities will require future investigations of the topological properties of the movie-driven activity.

The results of this study provide a functional anatomical framework for how deficits within the TPJ-pSTS may affect social cognition in schizophrenia. While the general ISC deficits replicate previous findings (27), the remaining results have not been described before and need to be replicated in a larger sample. One important limitation to note is that ISC results can be confounded by differences in visual scanning patterns, where group differences in which visual features are being fixated may drive differences in cortical activity. However, large differences were not observed in visual cortex, which would be most affected by differences in eye-gaze position. In addition, the increased correlation of the TPJ in SzP with HC prefrontal areas suggests that not *all* correlational results in SzP will be decreased by differences in gaze position. Another concern may stem from the ROI selection based on our own task localizers versus other available parcellation schemes. However, the grayordinate results, especially for the ISC, demonstrate that our results are likely robust to ROI selection biases. Lastly, these correlational results provide the necessary framework to test causality with TMS or other neuromodulation modalities. In summary, these results all converge on the TPJ-pSTS as functionally impaired in SzP, reducing its contribution to the transformation of visual information to a visual scanning plan and leading to social cognition deficits.

---

| Demographics/Performance<br>Mean (Standard Deviation) | Schizophrenia (MRI) (n=27) | Healthy Controls (MRI) (n=21) | Statistics (MRI) | Schizophrenia (Behavioral) (n=20) | Healthy Controls (Behavioral) (n=22) | Statistics (Behavioral) |
| --- | --- | --- | --- | --- | --- | --- |
| Age (years) | 38.6 (11.7) | 34.5 (9.6) | $t_{49}=1.30, p=0.20$ | 39.9 (11.7) | 34.6 (9.6) | $t_{42}=1.60, p=0.12$ |
| Gender (male/female) | 19/8 | 10/11 | $\chi^2=2.6, p=0.11$ | 13/7 | 12/10 | $\chi^2=0.48, p=0.49$ |
| Race/Ethnicity (%White %Black %Hispanic) | 33%<br>40.7%<br>25.9% | 38.1%<br>42.9%<br>14.3% | $\chi^2=0.11, p=0.73$<br>$\chi^2=0.02, p=0.88$<br>$\chi^2=0.97, p=0.32$ | 35.0%<br>40.0%<br>15.0% | 22.7%<br>59.1%<br>22.7% | $\chi^2=0.77, p=0.34$<br>$\chi^2=1.53, p=0.22$<br>$\chi^2=0.41, p=0.52$ |
| Participant Education (years) | 14.4 (3.0) | 15.2 (1.9) | $t_{49}=-0.11, p=0.31$ | 13.7 (2.4) | 15.1 (1.8) | $t_{42}=-1.97, p=0.06$ |
| Participant SES | 33.2 (14.4) | 38.2 (12.4) | $t_{49}=-1.04, p=0.21$ | 27 (10.7) | 34.3 (13.4) | $t_{42}=-2.0, p=0.06$ |
| Edinburgh Handedness Score | 17.3 (4.7) | 14.8 (6.5) | $t_{49}=1.38, p=0.18$ | 16.9 (3.4) | 16.8 (4.8) | $t_{42}=0.03, p=0.98$ |
| IQ (WRAT scaled score) | 97.3 (13.0) | 100.4 (14.6) | $t_{49}=-0.70, p=0.49$ | 97.2 (11.7) | 104.0 (11.9) | $t_{42}=-1.75, p=0.09$ |
| PANSS Positive Symptoms | 15.9 (5.5) | --- | --- | 16.5 (4.5) | --- | --- |
| PANSS Negative Symptoms | 13.5 (4.4) | --- | --- | 14.6 (4.1) | --- | --- |
| PANSS Total Scores | 58.4 (14.9) | --- | --- | 62.4 (15.3) | --- | --- |
| Antipsychotic dose (CPZ equivalents, mg) | 691.9 (1034.5) | --- | --- | 567.4 (496.5) | --- | --- |

## References

1. Green MF, Horan WP, Lee J (2015): Social cognition in schizophrenia. 16:620–631.

2. Ochsner KN (2008): The social-emotional processing stream: five core constructs and their translational potential for schizophrenia and beyond. In:. Biol Psychiatry. (Vol. 64), Presented at the Biological psychiatry, pp 48–61.

3. Corbetta M, Patel GH, Shulman GL (2008): The reorienting system of the human brain: from environment to theory of mind. Neuron. 58:306–324.

4. Polosecki P, Moeller S, Schweers N, Romanski LM, Tsao DY, Freiwald WA (2013): Faces in Motion: Selectivity of Macaque and Human Face Processing Areas for Dynamic Stimuli. J Neurosci. 33:11768–11773.

5. Koster-Hale J, Saxe R (2013): Theory of mind: a neural prediction problem. Neuron. 79:836–848.

6. Patel GH, Sestieri C, Corbetta M (2019): The evolution of the temporoparietal junction and posterior superior temporal sulcus. Cortex. doi: 10.1016/j.cortex.2019.01.026.

7. Power JD, Schlaggar BL, Lessov-Schlaggar CN, Petersen SE (2013): Evidence for hubs in human functional brain networks. Neuron. 79:798–813.

8. Russ BE, Leopold DA (2015): Functional MRI mapping of dynamic visual features during natural viewing in the macaque. 109:84–94.

9. White BJ, Berg DJ, Kan JY, Marino RA, Itti L, Munoz DP (2017): Superior colliculus neurons encode a visual saliency map during free viewing of natural dynamic video. Nat Comms. 8:14263.

10. Kay SR, Fiszbein A, Opler LA (1987): The positive and negative syndrome scale (PANSS) for schizophrenia. Schizophr Bull. 13:261–276.

11. First MB, Spitzer RL, Gibbon M, Williams JBW (1997): Structured Clinical Interview for DSM-IV® Axis I Disorders (SCID-I), Clinician Version, Administration Booklet. American Psychiatric Publishing, Inc.

12. McDonald S, Flanagan S, Rollins J, Kinch J (2003): TASIT: A new clinical tool for assessing social perception after traumatic brain injury. J Head Trauma Rehabil. 18:219–238.

13. Patel GH, Arkin SC, Ruiz-Betancourt D, DeBaun H, Strauss NE, Bartel LP, et al. (n.d.): What you see is what you get: visual scanning failures of naturalistic social scenes in schizophrenia. Psychol Med.

14. Glasser MF, Sotiropoulos SN, Wilson JA, Coalson TS, Fischl B, Andersson JLR, et al. (2013): The minimal preprocessing pipelines for the Human Connectome Project. doi: 10.1016/j.neuroimage.2013.04.127.

15. Power JD, Mitra A, Laumann TO, Snyder AZ, Schlaggar BL, Petersen SE (2014): Methods to detect, characterize, and remove motion artifact in resting state fMRI. 84:320–341.

16. Simony E, Honey CJ, Chen J, Lositsky O, Yeshurun Y, Wiesel A, Hasson U (2016): Dynamic reconfiguration of the default mode network during narrative comprehension. Nat Comms. 7:12141.

17. Grinband J, Steffener J, Razlighi QR, Stern Y (2017): BOLD neurovascular coupling does not change significantly with normal aging. Hum Brain Mapp. doi: 10.1002/hbm.23608.

18. Woolrich MW, Behrens TEJ, Beckmann CF, Jenkinson M, Smith SM (2004): Multilevel linear modelling for FMRI group analysis using Bayesian inference. 21:1732–1747.

19. Winkler AM, Ridgway GR, Webster MA, Smith SM, Nichols TE (2014): Permutation inference for the general linear model. 92:381–397.

20. Horiguchi H, Wandell BA, Winawer J (2016): A Predominantly Visual Subdivision of The Right Temporo-Parietal Junction (vTPJ). 26:639–646.

21. Martinez A, Gaspar PA, Hillyard SA, Andersen SK, Lopez-Calderon J, Corcoran CM, Javitt DC (2018): Impaired Motion Processing in Schizophrenia and the Attenuated Psychosis Syndrome: Etiological and Clinical Implications. American Journal of Psychiatry. appiajp201818010072.

22. Chen Y, Palafox GP, Nakayama K, Levy DL, Matthysse S, Holzman PS (1999): Motion perception in schizophrenia. Arch Gen Psychiatry. 56:149–154.

23. Martinez A, Tobe R, Dias EC, Ardekani BA, Veenstra-VanderWeele J, Patel GH, et al. (2019): Differential Patterns of Visual Sensory Alteration Underlying Face Emotion Recognition Impairment and Motion Perception Deficits in Schizophrenia and Autism Spectrum Disorder. Biol Psychiatry. doi: 10.1016/j.biopsych.2019.05.016.

24. Chen Y, Levy DL, Sheremata S, Holzman PS (2004): Compromised late-stage motion processing in schizophrenia. Biol Psychiatry. 55:834–841.

25. Kohler CG, Walker JB, Martin EA, Healey KM, Moberg PJ (2010): Facial Emotion Perception in Schizophrenia: A Meta-analytic Review. Schizophr Bull. 36:1009–1019.

26. Taylor SF, Kang J, Brege IS, Tso IF, Hosanagar A, Johnson TD (2012): Meta-analysis of functional neuroimaging studies of emotion perception and experience in schizophrenia. Biol Psychiatry. 71:136–145.

27. Mäntylä T, Nummenmaa L, Rikandi E, Lindgren M, Kieseppä T, Hari R, et al. (2018): Aberrant Cortical Integration in First-Episode Psychosis During Natural Audiovisual Processing. Biol Psychiatry. doi: 10.1016/j.biopsych.2018.04.014.

28. Yeo BTT, Krienen FM, Sepulcre J, Sabuncu MR, Lashkari D, Hollinshead M, et al. (2011): The organization of the human cerebral cortex estimated by intrinsic functional connectivity. J Neurophysiol. 106:1125–1165.

29. Croxson PL, Forkel SJ, Cerliani L, Thiebaut de Schotten M (2017): Structural Variability Across the Primate Brain: A Cross-Species Comparison. 1–13.

30. van Essen DC, Dierker DL (2007): Surface-based and probabilistic atlases of primate cerebral cortex. Neuron. 56:209–225.

31. Power JD, Cohen AL, Nelson SM, Wig GS, Barnes KA, Church JA, et al. (2011): Functional network organization of the human brain. Neuron. 72:665–678.

32. Gordon EM, Laumann TO, Adeyemo B, Huckins JF, Kelley WM, Petersen SE (2014): Generation and Evaluation of a Cortical Area Parcellation from Resting-State Correlations. doi: 10.1093/cercor/bhu239.

33. Harvey P-O, Zaki J, Lee J, Ochsner KN, Green MF (2013): Neural substrates of empathic accuracy in people with schizophrenia. Schizophr Bull. 39:617–628.

34. Lee J, Zaki J, Harvey P-O, Ochsner KN, Green MF (2011): Schizophrenia patients are impaired in empathic accuracy. Psychol Med. 41:2297–2304.

35. Zaki J, Hennigan K, Weber J, Ochsner KN (2010): Social cognitive conflict resolution: contributions of domain-general and domain-specific neural systems. J Neurosci. 30:8481–8488.

36. Andrews-Hanna JR, Reidler JS, Huang C, Buckner RL (2010): Evidence for the default network’s role in spontaneous cognition. J Neurophysiol. 104:322–335.

37. Whitfield-Gabrieli S, Thermenos HW, Milanovic S, Tsuang MT, Faraone SV, McCarley RW, et al. (2009): Hyperactivity and hyperconnectivity of the default network in schizophrenia and in first-degree relatives of persons with schizophrenia. Proc Natl Acad Sci USA. 106:1279–1284.

38. Baker JT, Holmes AJ, Masters GA, Yeo BTT, Krienen F, Buckner RL, Ongür D (2013): Disruption of Cortical Association Networks in Schizophrenia and Psychotic Bipolar Disorder. JAMA Psychiatry. doi: 10.1001/jamapsychiatry.2013.3469.

39. Alexander-Bloch AF, Vértes PE, Stidd R, Lalonde F, Clasen L, Rapoport J, et al. (2013): The anatomical distance of functional connections predicts brain network topology in health and schizophrenia. Cereb Cortex. 23:127–138.

