## Supplemental Materials for "Failure to engage the TPJ-pSTS during naturalistic scene processing in schizophrenia"

### Supplementary Methods

*Behavioral Movie Eye-tracking and Social Cognition Assessment*

To measure eye-tracking performance during the free-viewing of a naturalistic scene, we measured eye-movements while participants watched the first 15 minutes of the cinematic movie "The Good, the Bad, and the Ugly" (United Artists, 1966) with the audio track silenced. Participants were seated 72cm from a 27" monitor (BenQ XL2720Z, BenQ USA, Costa Mesa CA), with their heads resting comfortably in a chin/head rest. The size of the movie was 15.7° x 8.2° visual angle. An eye-tracker (EyeLink 1000plus, SR Research, Mississauga, Ontario, Canada) recorded eye-movements at 1000Hz. Participants were asked to briefly summarize the clip afterwards to ensure attention.

To measure social cognition, participants viewed the 32 video clips (23.0° x 19.1°) in TASIT Part 3. TASIT Part 3 consists of an A and B section. The B section contains the same scenarios with the same dialogue as A, but with the sarcasm/lie instructions flipped. After each video clip, a research assistant verbally asked participants 4 questions about what the main character was thinking, feeling, doing, and saying, and recorded their responses with no time limit restriction. Only TASIT 3-A data were considered in this study due to the possibility of memory effects on TASIT 3-B performance.

*Apparatus and Procedural Setup for fMRI*

Participants were placed on the scanner bed with cushioning to stabilize potential head movement and earplugs to block out MR-related sound. A mirror attached to the head coil allowed participants to view visual stimuli displayed on a translucent screen mounted at the end of the bore from a liquid crystal display (LCD) projector; the eye-to-screen distance (mirror-to-screen + eye-to-mirror) was 107 cm. Participants held a fiber optic, five-channel button box (Current Designs, Philadelphia, PA) in their right hands that allowed them to respond to target stimuli with their index finger. The experimental display was controlled by a MacBook Pro (Apple Inc., Cupertino, CA) running custom software within either MATLAB (Mathworks, Natick, MA) using Psychophysics toolbox version 3.012 (Brainard, 1997; Pelli, 1997) or Experiment Builder (SR Research, Mississauga, Ontario, Canada), which connected to the projector via high-definition multimedia interface (HDMI) hub. The stimuli were synchronized to the beginning of the MR volume acquisition via a USB device that triggered the MATLAB paradigm to begin (Current Designs, Philadelphia, PA). Eye movements were recorded using Eyelink 1000 plus (SR Research, Mississauga, Ontario, Canada).

*fMRI Acquisition*

Functional and anatomical data were acquired with a 32-channel phased array receive-only head coil (Nova Medical, Wilmington, MA) by General Electric's Discovery MR750 3.0 Telsa full body MR scanner (GE; Fairfield, CT) at New York Psychiatric Institutes' (NYSPI) MRI Research Unit. Localizer scans and a gradient echo B0 fieldmap (TE=4.8ms, TR=800ms, FOV=256mmx256mm, slice thickness=2mm, matrix=128x128, slices=68) were required for the Human Connectome Project (HCP) processing pipelines. Structural images were acquired in 5-minute sequences and were comprised of two T1-weighted images (3D sagittal, 0.8mm isotropic, matrix size=300x300, slices=220, TR=7856ms, TE=3108ms, flip angle=12, TI=450ms) and two T2-weighted images (3D sagittal, 0.8 mm isotropic, matrix size=320x320, slices=220, TR=2500ms, TE=95.708, flip angle=90°). Task and resting state functional data were collected with a multiband SMS-EPI sequence (2 mm isotopic, slice plane=transverse, TR=850ms, MUX=6, ARC=1, TE=25ms, matrix size=96x96, slices=66, phase encoding direction=P→A) (courtesy of the Center for Cognitive and Neurobiological Imaging, Stanford University, http://cni.stanford.edu).

*Movie-watching Task*

Participants viewed the same 15 minute clip of the movie "The Good, the Bad, and the Ugly" as above with the audio track silenced while eye-movements were recorded. Participants were asked to briefly summarize the clip afterwards to ensure attention. BOLD-fMRI data were collected in a single run of 1068 frames.

*Attention Localizer*

Attention areas were localized with an RSVP visual search task similar to the one used by Patel *et al.* {Patel:2015cf}{Patel:2010hz}. 34 HCs performed this task. Forty-two clipart images of neutral objects were used as both target and distractor stimuli. One of the forty-two clipart images was randomly selected as the

target image, while the other forty-one images were used as distractors. Before the BOLD run started, the target image was presented once in each of the three stream locations to familiarize the participant with the target. One stream was superimposed over the fixation point in the center of the screen with a width/height of 3.6□. The two peripheral streams were located at 5.8□ eccentricity with a polar angle of 0□ (left of the fixation point) and 180□(right of the fixation point) and with a width/height of 5.3□. Targets were chosen randomly, under the condition that no target could be used twice in a row. Participants pressed a specified button on a button box with their right index finger every time they saw the target stimulus.
Each trial consisted of a 10 second RSVP stream in one of three locations and with the same sizes described above. Each stimulus in the RSVP stream was displayed for 100 milliseconds (ms) before being replaced by the next distractor with no gap between presentations. Distractor stimuli were chosen randomly with the constraint of not having consecutive repeats. The target stimulus took the place of a distractor stimulus in the stream and did not differ in any parameters from the distracting stimuli. Half of all trials had a single target presented randomly within the first 8 seconds in the 10-second stream. One-quarter of trials had no target (catch trials). In the remaining one-quarter of trials, the target appeared twice within the target stream, once within the first 4 seconds and again from 4 to 8 seconds. After the target appeared, participants had 1.3 seconds to indicate detection. Participants were instructed to respond as accurately and quickly as possible, with an emphasis on accuracy so as to minimize false positives. Following each 10-second stream, the participant was asked to maintain fixation for inter-trial intervals of 2, 4, or 6 seconds, chosen randomly, before the next trial began.  Two BOLD-fMRI runs were collected, each 5 minutes 30 seconds each (~20 trials / run).

*Static/Moving Emotional Faces Localizer*
To localizes areas involved in the processing of static and moving facial expressions, participants passively viewed blocks of static or moving emotional or neutral stimuli.  The four block types were: 1) “neutral static”, five pictures of the faces of differing people displaying no emotion; 2) “neutral moving”, five videos of the faces of differing people moving but displaying no emotion; 3) “emotion static”, five pictures of the faces of differing people displaying a certain emotion (happiness, anger, fear, sadness);  4) “emotion moving”: five videos of the faces of differing people displaying the same emotions.  All stimuli were in grayscale.  Pictures were displayed for 2 seconds each, and videos for 4 seconds each, with no gap between either.  Blocks were repeated in this order, and interspersed with 20s of fixation.  Two runs of 7 minutes 5 seconds each were collected from 24 HC.

*Motion Localizer*
To identify the motion-sensitive visual areas, we used a previously described motion localizer {Tootell:1995uc}{Martinez:2018be}. In short, participants maintained fixation on a central cross while a series of low-contrast concentric rings of a diameter of 15 degrees expanded or contracted for 20 seconds.  These blocks alternated with 20 second blocks during which the same rings remained static. Participants were instructed to respond to occasional dimmings of the fixation point via a button press to maintain attention and fixation.  One run of 5 minutes 23 seconds each were collected from 14 HC.

*Mentalization (Theory of Mind) Localizer*
To identify areas involved in mentalization, we used a previously described movie-watching task {Jacoby:2016co}.  Participants were instructed to watch the short, animated clip of Disney Pixar’s “Partly Cloudy”, and then briefly summarize the plot afterwards to ensure attention.  A single run of 6 minutes 36 seconds was collected for 12 HCs.

*Image Processing*
All imaging data were processed on workstation machines (Mac Pro, Apple Inc., Cupertino, CA) using The HCP processing pipeline v3.4 adapted for the NYSPI GE MR750 MRI scanner. The HCP pipelines first processed the anatomy images to create a cortical surface model for each individual aligned to the HCP fs_LR 32k atlas, then for the functional runs it performed movement correction, distortion correction, and atlas alignment in a single resampling step, and lastly projected the functional data to an atlas cortical surface through the individual’s cortical surface model (Glasser et al., 2013). After each step, manual quality checks were performed to check for errors in atlas-alignment and segmentation. This pipeline is different from other processing pipelines in that it creates a Connectivity Informatics Technology Initiative (CIFTI) file for each

BOLD run that only contains the data from the cortical and the subcortical grey matter (“gray-ordinates” as opposed to voxels), which allows for a more precise localization of brain activity that is not confounded by cerebrospinal fluid (CSF) or white matter partial volume effects. Structural and functional images were aligned in a volume space of the Montreal Neurological Institute (MNI152) atlas and on the surface Conte69/fs_LR 32k atlas created by the HCP pipeline developers (Evans et al., 1992; Glasser et al., 2013; van Essen et al., 2012).

Additional post-processing procedures, adapted from Power et al. (Power et al., 2014), were performed to further minimize noise. First, estimates of head motion calculated in the X, Y, and Z directions along with the displacements of rotation around the X, Y, and Z axes (pitch, yaw, and roll) by the HCP movement correction algorithm, along with their derivatives (backwards difference), the square of the six parameters, and the sum of these six parameters (assuming a 50mm radius of head size for the rotations, previously labeled as the framewise displacement (FD)) were used as nuisance regressors to remove movement-related artifact. Second, tissue signals and their derivatives calculated by averaging signal across voxels within a spatial mask for ventricle signals, CSF signal, white matter signal, and whole brain signal were used as nuisance regressors to remove physiology-related noise (global signal regression also removes movement-related artifact (Power, 2016). Third, MR frames with an FD greater than the 75%tile + 1/2 * interquartile range of the FD trace for each run, limited to a range between 0.2mm and 0.5mm were censored, and replaced by interpolation using a method based on the Lomb-Scargle periodogram (Lomb, 1976; Power et al., 2014). Fourth, data were low band-pass filtered at .588 Hz (the Nyquist limit for the MR sampling rate) and high band-pass filtered at 0.0005 Hz (Glasser et al., 2013). The resulting time-series was used for all subsequent analyses. All analyses were repeated with the FD threshold set to 0.2mm and with no whole brain signal regression.

*Task fMRI Analyses*

The task fMRI analyses for the attention, motion, and static/moving faces paradigms were all conducted using FEAT {Woolrich:2004it}. For level 1 analyses, the magnitudes of activation evoked by attention to/processing of paradigm-related “events” were separately estimated in each individual via a general linear model (GLM) by modeling each event type as an independent regressor {Shulman:2003gc}. For each paradigm, the regressor was formed by convolving a boxcar function corresponding to the event length with an estimate of the hemodynamic response to an impulse function {Grinband:2017cz}. A high-pass filter was of the length of the BOLD run was used to remove linear trend. The first four volumes of each run were excluded from the analyses. Results from individuals with two BOLD runs were combined in a level 2 fixed-effects analysis. Level 3 group analyses were performed using FLAME 1 mixed-effects.

For the attention localizer, the magnitudes of activation evoked by attention to and processing of the RSVP stream and detection of the target at each spatial location were separately estimated in each individual. For the RSVP stream, the event duration was 10 seconds. For detection events, the reaction time of the response to each target was used as the event duration. For the misses and false detections, the event duration used was 500ms. For the static/moving faces localizer, the modeled event blocks were neutral moving, neutral static, emotion static, and emotion moving. The event durations used for convolution were 20 seconds for the moving and 10 seconds for the static. For the motion localizer, the only modeled event was for the blocks of the expanding/contracting rings, which had a length of 12 seconds.

To analyze the mentalization localizer, we utilized the hand-coded mental, social, physical pain and control regressors for the animated short as described in Jacoby *et al.* {Jacoby:2016co}. The four regressors were then convolved with the same estimate of the hemodynamic response to an impulse function used above {Grinband:2017cz}. A one-sample t-test was performed in MATLAB (Natick, MA) to create the statistical map of activations.

*ROI Definitions*

For each group, regions of interest (ROIs) were defined on the average activation maps on the cortical surface by drawing borders around each activation. Activation gradient maps derived from the spatial derivative of the z-statistic maps were used to define borders around local maxima (peaks), and to separate contiguous activations by tracing the line that represented the highest gradient (local minimum) between the two activations (Glasser et al., 2016). ROI labels were largely based on sulcal/gyral anatomy, with the HCP’s multimodal parcellation of the cortical surface serving as a guide for labeling some of the ROIs. Foci spanning several HCP ROIs were labeled according to the main overlapping ROI (Glasser et al., 2016). TPJa was drawn on the mixed-effects maps of the conjunction of gray-ordinates a) deactivated by the RSVP monitoring and b) activated by target detection (Patel et al., 2015), and was limited to the posterior bank of the supramarginal gyrus.

*Intersubject Correlation Analysis*
Inter-participant correlation (ISC) values of each voxel for each participant (HC n=21; SzP n=27) were calculated as the average pairwise correlation between the time-course of the BOLD signal from each grayordinate from each participant evoked by the movie and the time-course of the BOLD signal evoked by the same stimulus of the same grayordinate in all other HC participants. SzP participants were correlated with all HC participants; all HC participants were correlated with all HC participants except themselves. These average correlation values were Fisher-z transformed.  For ROI ISC, analyses, the same procedure was followed except time-courses came from each ROI, not each grayordinate.

*TPJ Synchronization Analysis*
For each ROI, the average time-course of activity was extracted for the HCs.  This time-course was then correlated to the time-course of activity for each ROI or grayordinate in each individual for each group. Individual HCs were correlated against the average of all other HCs.  The individual ROI correlations were then Fisher-z transformed for group-level statistics.

*Visual Feature Correlation Analyses*
The average luminance, contrast, and motion energy of all pixels were calculated for each video frame of the movie, resulting in three separate visual feature regressors {Russ:2015gh}.  A fourth regressor was also created representing the number of faces on screen on each video frame (http://aws.amazon.com/rekognition). All three video regressor time series were down-sampled to match the acquisition frequency of the corresponding BOLD signal and then convolved with an approximate hemodynamic response function (HRF) {Grinband:2017cz}.  The motion regressor was then log-transformed for normality; contrast and luminance were already normally distributed.  Each video regressor was then independently correlated with the BOLD signal induced by the GBU stimulus of each ROI or grayordinate of each participant. These correlation values were then Fisher-z transformed.

*Saccade Frequency Correlation Analysis*
A saccade frequency regressor was calculated for each participant with useable eye-tracking data as the number of saccades made by that participant during the 850 ms sampling interval (TR). This regressor was convolved with an approximated HRF {Grinband:2017cz}. The average number and average frequency of saccades were calculated for each group. A participant's saccade frequency regressor was correlated with the same participant's GBU stimulus-induced BOLD signal at all ROIs or grayordinates and then Fisher-z transformed.

**Supplementary Figures**

**Supplemental Figure 1:** The localizer task activations clearly separates grayordinates activated by static and moving facial expressions in the pSTS (purple borders) versus mentalization on the angular gyrus in the posterior TPJ (TPJp, yellow border) (**Supplemental Figure 1 F/G** versus **1B**). The localizers also helped to divide the TPJp from the anterior TPJ on the supramarginal gyrus (TPJa, green border) involved in visual search (deactivations in **Supplemental Figure 1C**) and detection (activations in **Supplemental Figure 1D**). In between these ROIs in the TPJ was an undefined section of cortex; this ROI was assigned the name TPJm (red border) .
