## Supplementary figures and images for "Failure to engage the TPJ-pSTS during naturalistic scene processing in schizophrenia"

### Supplemental Figure 1

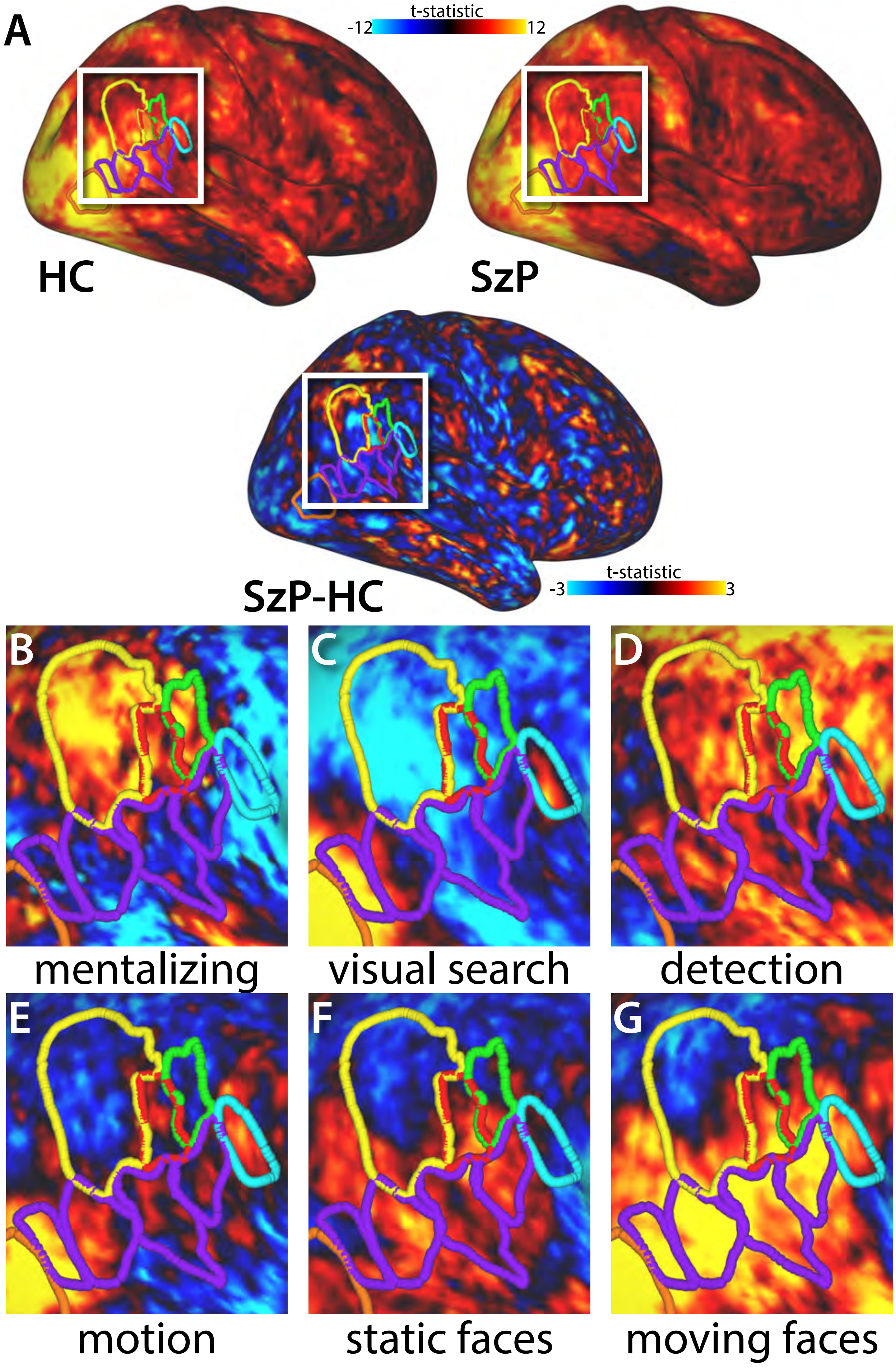
